# Hypoxia inducible factors inhibit respiratory syncytial virus infection by modulation of nucleolin expression

**DOI:** 10.1101/2023.10.31.564923

**Authors:** Xiaodong Zhuang, Giulia Gallo, Parul Sharma, Jiyeon Ha, Andrea Magri, Helene Borrmann, James M Harris, Senko Tsukuda, Eleanor Bentley, Adam Kirby, Simon de Neck, Hongbing Yang, Peter Balfe, Peter AC Wing, David Matthews, Adrian L Harris, Anja Kipar, James P Stewart, Dalan Bailey, Jane A McKeating

## Abstract

Respiratory syncytial virus (RSV) is a global healthcare problem, causing respiratory illness in young children and elderly individuals. However, our knowledge of the host pathways that define susceptibility to RSV infection and disease severity is limited. Hypoxia inducible factors (HIFs) define cellular metabolic processes in response to low oxygen and are recognised to play a key role in regulating inflammatory responses in the lower respiratory tract. We demonstrate a role for hypoxia inducible factors (HIFs) to suppress RSV entry and RNA replication. We show that hypoxia and HIF prolyl-hydroxylase inhibitors significantly reduce expression of the RSV entry receptor nucleolin and inhibit viral driven cell-cell fusion. We identify a HIF regulated microRNA, miR-494, that regulates nucleolin expression. In RSV-infected mice, treatment with the clinically approved HIF prolyl-hydroxylase inhibitor, Daprodustat, reduced the level of infectious virus and infiltrating monocytes and neutrophils in the lung. This study highlights a role for HIF-signalling to limit multiple aspects of RSV infection and associated inflammation and informs future therapeutic approaches for treating this respiratory pathogen.

## INTRODUCTION

Respiratory syncytial virus (RSV) causes lower respiratory tract illness in young children^1^ and is increasingly recognised as a significant respiratory pathogen in immunocompromised adults and the elderly^2^. The virus is primarily transmitted through close contact, although it can spread via aerosolized droplets^3^. Following a brief period of replication in the epithelium of the nasopharynx and upper respiratory tract, RSV can infect the small bronchioles or alveoli of the lower respiratory tract^4^. Treatment options are limited and the humanized monoclonal antibody (Palivizumab) targeting the viral encoded fusion (F) protein is only used for passive immunoprophylaxis of high-risk infants^5^. The recent FDA approval of RSV vaccines for immunizing elderly populations^6^ provides opportunities to reduce the burden of RSV associated disease and highlights the continued need to develop new anti-viral therapies for this infection.

During severe RSV infection, the host’s immune response triggers an increase in mucus production and inflammation, leading to airway narrowing, bronchiolitis in young children and acute respiratory illness in the elderly or those with underlying chronic conditions^7^. Virus induced damage to the alveolar epithelia can reduce local oxygen levels and is a common feature of RSV bronchiolitis that can result in respiratory failure, apnea, and death^8,9^. Hypoxia inducible factors (HIFs) regulate the cellular transcriptome and metabolome in response to low oxygen and inflammatory conditions^10^. HIFα subunits are regulated by oxygen-dependent pathways and when oxygen is abundant, newly synthesised proteins are hydroxylated by HIF prolyl-hydroxylase domain (PHD) enzymes and targeted for proteasomal degradation. When oxygen is limited the PHD enzymes are inactive and HIFα dimerizes with HIF-1β to activate transcription of genes regulating cell metabolism, pulmonary vasomotor control and immune regulation^11–13^. HIF regulated genes can vary between cell types allowing a flexible and tissue specific response to diverse physiological signals^14^.

HIFs have been reported to enhance endogenous pathways to protect against lung injury during acute inflammation that include: promoting vascular repair^15^; suppressing the inflammatory response in bronchial epithelial cells and reducing interleukin 6 and interferon gamma-induced protein 10 expression^16^; regenerating alveolar type II pneumocytes^17^; preventing the bioenergetic failure of alveolar epithelial cells during Acute Respiratory Distress Syndrome (ARDS)^18–20^ and interacting with key metabolites such as succinate to limit damage from acute lung injury^21^. Zhao et al reported that lung epithelial *Hif1a* knockout mice support higher levels of influenza A virus infection and associated mortality^22^. We previously reported that hypoxic or pharmacological activation of HIFs limit SARS-CoV-2 infection and epithelial damage in the Syrian Golden Hamster model of COVID-19^23,24^. Collectively, these studies highlight the significance of HIFs in defining host susceptibility to respiratory viruses and associated pathologies. Importantly this has the potential for pharmacological intervention since drugs inhibiting the PHD enzymes that drive HIF-mediated erythropoiesis are licensed for the treatment of renal anaemia^25–30^. These studies implicate HIF activation as a therapeutic strategy to combat lung injury.

RSV is primarily cell-associated and the virus can spread by cell-cell fusion, generating syncytia - large multinucleated cells formed by the fusion of infected with neighbouring uninfected cells. The RSV fusion protein (RSV-F) has important roles in entry and fusion, driving particle attachment to cellular receptors and fusion of virion and host-cell membranes, either at the cell surface or during particle macropinocytosis^31^. Several host factors including intercellular adhesion molecule 1 (ICAM1)^32^, epidermal growth factor receptor (EGFR)^33^ and nucleolin (NCL)^34^ have been shown to facilitate RSV mediated cell-cell fusion. In our study, we show that hypoxic conditions or treatment with PHD inhibitors (PHI) limits RSV infection via perturbing expression of nucleolin via the hypoxia-induced microRNA, miR-494. Furthermore treatment of RSV infected mice with the clinically approved PHI, Daprodustat, reduced the level of infectious virus and infiltrating monocytes and neutrophils in the lung, highlighting a role for HIF-signalling in regulating host responses to respiratory viruses.

## RESULTS

### Hypoxia and HIF prolyl-hydroxylase inhibitors limit RSV infection

To investigate a role for HIFs in regulating RSV replication we cultured human lung epithelial Calu-3 cells at 18% or 1% oxygen for 24h before infecting with RSV (A2 strain, MOI 0.2). Infection was quantified by measuring intracellular viral RNAs encoding fusion (F) and infectious virus at 24h and 48h post-infection (pi). We selected 1% oxygen as this is known to stabilise HIFα subunits and activate HIF target gene transcription. We confirmed the infected cells responded to hypoxia by immunoblotting for HIF-1α and showed increased expression of the HIF regulated N-myc downregulated gene (NDRG1) expression (**Fig.1a**). Hypoxic conditions significantly reduced the level of RSV-F transcripts and infectious virus at 48hpi. Earlier studies showed that RSV infection stabilised HIF-1α expression^35–37^. To investigate whether RSV induced a functional HIF response in our model system, we used Calu-3 expressing a hypoxia-response element (HRE) driven GFP reporter. Treating the reporter cells with three PHIs (Daprodustat, Molidustat and Roxadustat) induced GFP expression in a time-dependent manner, whereas infection with RSV failed to activate GFP expression or to stablise HIF-1α expression (**Fig.1b**). Furthermore, flow cytometric analysis to identify infected cells showed no evidence of increased GFP expression in the RSV-F expressing cells, demonstrating that infection does not activate HIF signalling in Calu-3 cells. We confirmed the majority of Calu-3 cells (>70%) expressed GFP following Daprodustat treatment demonstrating the cells were responsive to PHI treatment (**Fig.1c**).

**Figure 1.**
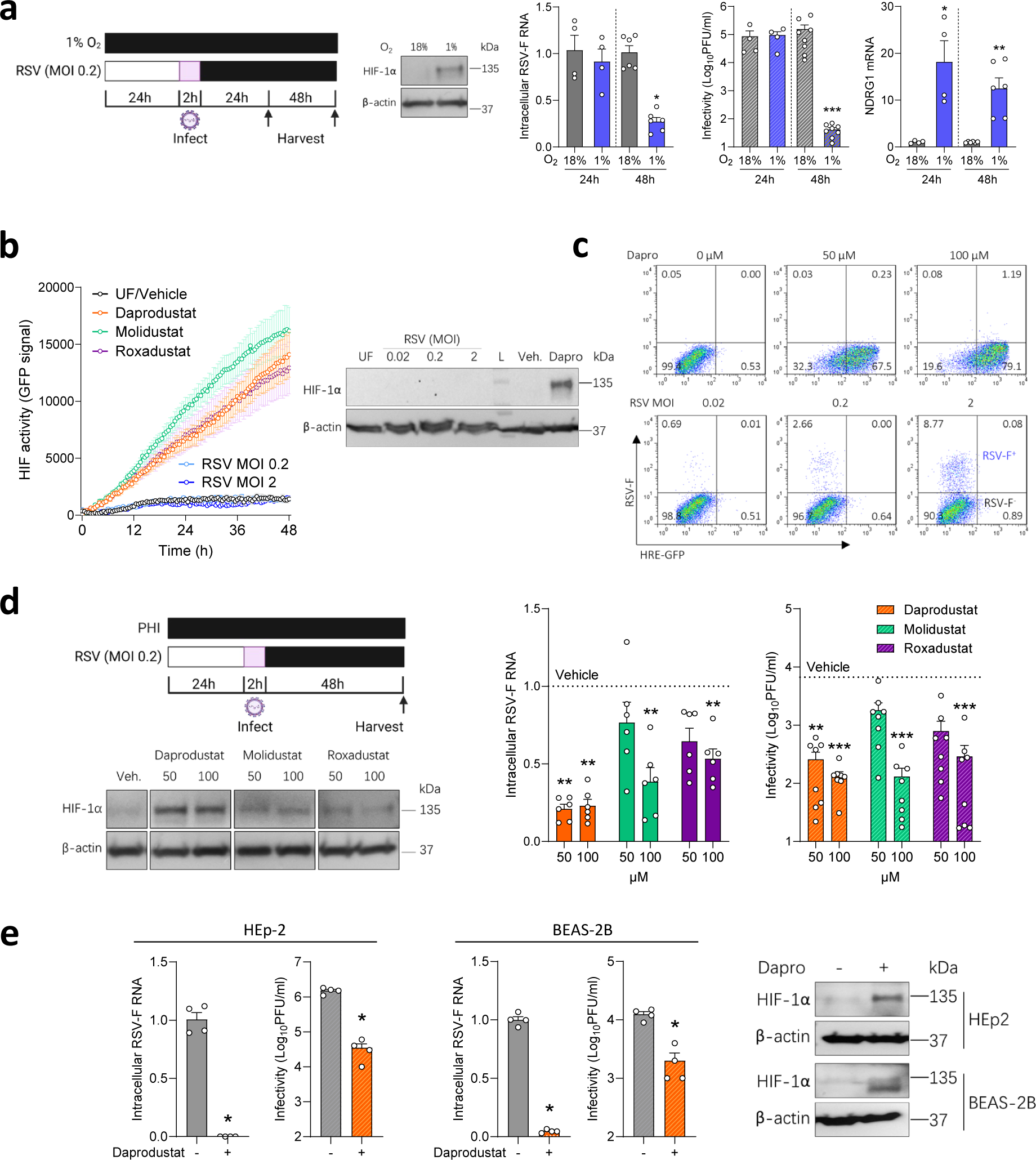
RSV infection is suppressed under hypoxic conditions and treatment with prolyl-hydroxylase inhibitors. **a**. Calu-3 cells were cultured at 18% (grey) or 1% (blue) O_2_ oxygen for 24h prior to infecting with RSV (MOI 0.2) and the cultures maintained under normoxic or hypoxic conditions for the stated times and cells lysed for PCR quantification of intracellular viral RNA (RSV-F), NDRG1 mRNA and infectivity at 24h or 48hpi (mean ± SEM, *n* = 4-8, Mann–Whitney test, Two-sided). All data are expressed relative to 18% O_2_ values at their respective time point. HIF-1α and β-actin protein expression were assessed by immunoblot at 48hpi. **b**. Real-time monitoring of Calu-3 cells transduced with HRE-GFP were treated with PHIs (Daprodustat, Molidustat or Roxadustat) or infected with RSV at MOI 0.2 or 2 (mean ± SEM, *n* = 5), where UF = uninfected. **c**. Calu-3 HRE-GFP reporter cells were treated with Daprodustat at 50 or 100 μM or infected with RSV at a range of MOIs for 48hpi and stained with anti-RSV-F-APC followed by flow cytometric analysis. **d**. Calu-3 cells were pre-treated with PHIs for 24h before infecting with RSV (MOI 0.2) and intracellular viral RNA (RSV-F) and infectivity measured at 48hpi (mean ± SEM, *n* = 6-8, One-way ANOVA with multiple comparisons, Two-sided). **e**. HEp-2 or BEAS-2B were pre-treated with Daprodustat (100 μM) for 24h before infecting with RSV (MOI 0.2) and intracellular viral RNA (RSV-F) and infectivity measured at 48 hpi (mean ± SEM, *n* = 4, Mann–Whitney test, Two-sided). HIF-1α and β-actin protein expression were assessed by immunoblot at 48hpi. UF = uninfected.

To ascertain a role for HIFs in regulating RSV infection, we treated Calu-3 cells with PHIs for 24h prior to infecting with RSV and demonstrated reduced levels of RSV-F transcript and infectious virus at 48hpi with no discernable effect on cell viability (**Fig.1d, Supplementary Fig.1**). HIF-1α was detected in the PHI treated cells, with Daprodustat inducing the highest expression (**Fig.1d**). Kinetic analysis of Daprodustat treated Calu-3 cells showed a time-dependent expression of HIF-1α and induction of NDRG1 gene expression at 24-48h post-treatment (**Supplementary Fig.2**). To assess whether Daprodustat limits RSV replication/transcription we used a transient RSV replicon assay encoding a luciferase reporter^38^. RSV-driven luciferase activity was dependent on the viral encoded polymerase and Daprodustat reduced RSV replication (**Supplementary Fig.3**). Viral RNA synthesis occurs in cytoplasmic inclusion bodies (IBs) that represent condensates of the viral replicase components formed by liquid-phase separation^39^. The steroidal alkaloid Cyclopamine was recently identified to target these IBs and limit viral replication and we demonstrated a potent inhibition of RSV-luciferase activity, consistent with the genesis of IBs in the RSV replicon assay (**Supplementary Fig.3**). The antiviral activity of Daprodustat was replicated in two additional cell lines (HEp-2 and BEAS-2B) that are commonly used in RSV research ^40^ and we noted a reduction in intracellular RSV-F transcripts and infectious virus (**Fig.1e**). Probing the Daprodustat or vehicle treated RSV infected cell lysates for HIF-1α confirmed expression in the drug treated cells (**Fig.1e**). We extended these observations to differentiated human primary bronchial epithelial cells (PBEC) propagated under air-liquid interface (ALI) and showed that Daprodustat reduced the level of infectious RSV secreted from the apical surface of these cultures (**Supplementary Fig.4**). We only detected HIF-1α expression or increased NDRG1 gene expression in the drug-treated ALI-PBEC, suggesting minimal evidence for RSV infection to activate HIFs in these model systems. Our previous experience of working with ALI-PBECs show limited HIF-1α expression under laboratory ‘normoxic’ conditions of 18% oxygen^23^. Collectively, these data show a role for hypoxia to limit RSV infection and pharmacological activation of HIF in multiple cell lines and PBEC, demonstrating its role in restricting RSV infection.

### Prolyl-hydroxylase inhibitors inhibit RSV replication kinetics

To investigate how HIFs regulate RSV replication, we used an RSV reporter system with GFP inserted as a separate transcription unit in the viral genome, allowing us to monitor viral replication kinetics. Calu-3 cells were pretreated with Daprodustat, Molidustat, or Roxadustat (50 μM) for 24h before infecting with the recombinant RSV (MOI 0.2). All three drugs inhibit RSV replication, with Daprodustat showing the most potent activity and screening a range of drug concentrations showed an IC_50_ of 9.0 µM at 48hpi (**Fig.2a** and **Supplementary Fig.5**). Bovine RSV (bRSV) is genetically closely related to human RSV, and we observed a consistent inhibition of bRSV replication by all three drugs (**Supplementary Fig.6**), highlighting a conserved interplay of the HIF signalling pathway across the orthopneumoviruses. We examined whether treating RSV infected Calu-3 cells (at 24 hpi or 48 hpi) with Daprodustat or Molidustat (50 μM) could influence virus replication and noted a significant suppression under these conditions (**Fig.2b**). These data are consistent with a role for the PHIs to limit virus transmission. Importantly, pre-treating Calu-3 cells with Daprodustat (used at an IC_50_ value of 9.0 μM) and Palivizumab, a clinically approved RSV antibody ^41,42^, showed greater antiviral activity of the drug combination than either treatment alone (**Fig.2c**). These data show that PHIs limit the kinetics of human and bovine RSV replication and a combined treatment of Daprodustat and Palivizumab shows additive antiviral activity.

**Figure 2.**
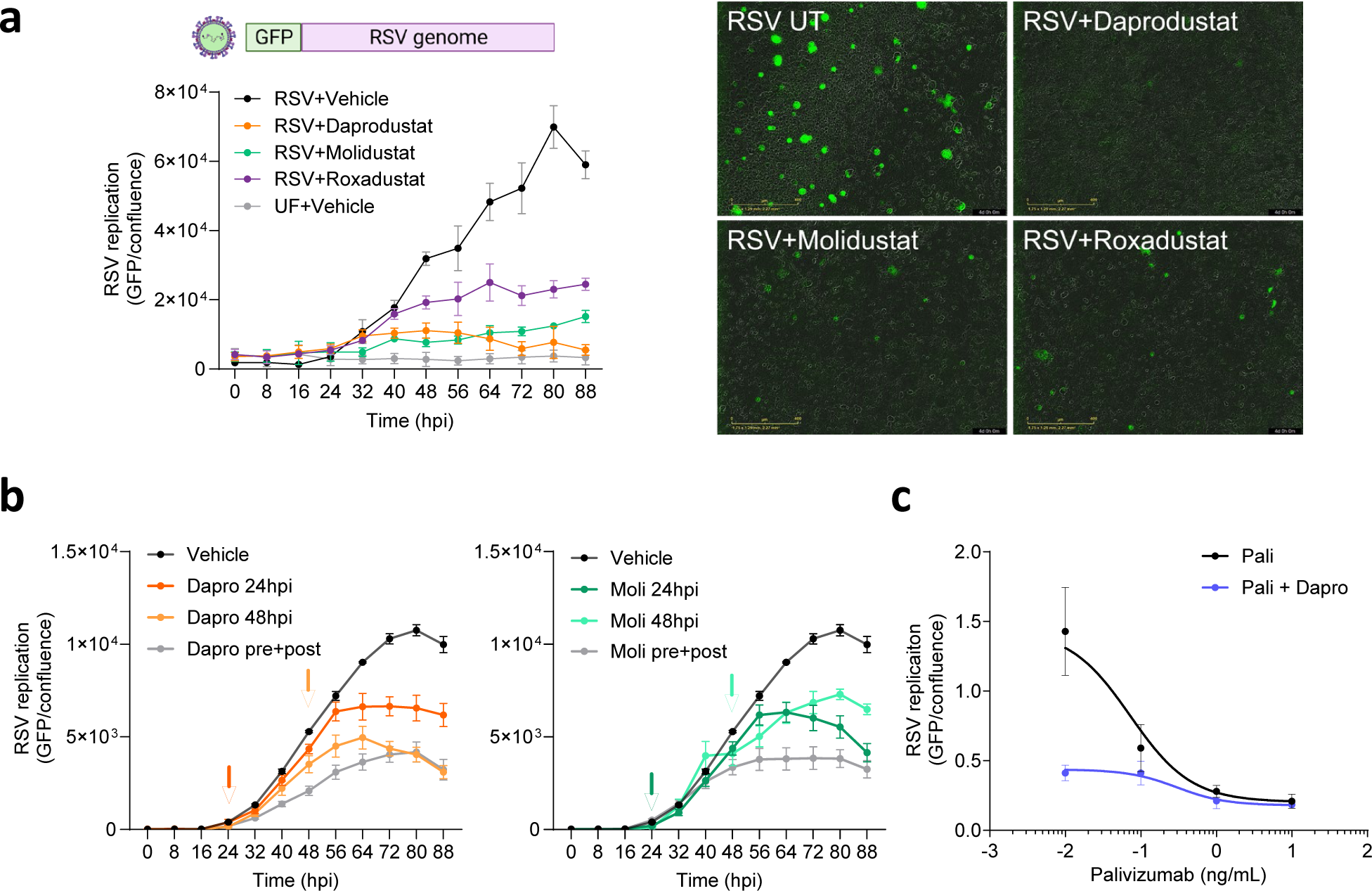
Prolyl-hydroxylase inhibitors inhibit RSV replication. **a.** Calu-3 cells were pre-treated with PHIs (Daprodustat, Molidustat, Roxadustat) 24h prior to infection with a reporter RSV expressing GFP (MOI 0.2) and viral replication measured every 8h using an Incucyte® Live-Cell Imaging System. GFP signal was normalised per image with respect to cell confluence (mean ± SEM, n = 4). **b**. Effect of PHI treatment on RSV infected Calu-3 cells. Calu-3 cells were treated with Daprodustat or Molidustat (50 μM) before and after infection (pre+post) or 24hpi or 48hpi where the arrows indicate addition of drugs and GFP expression measured using the Incucyte® Live-Cell Imaging System. GFP signal and confluence were measured and the integrated intensity values are represented (mean ± SEM, n = 6). **c.** Combined Daprodustat and Palivizumab treatment inhibits RSV replication. Calu-3 cells were treated with a range of Palivizumab doses in the presence or absence of Daprodustat (9.0 μM) prior to infection with RSV-GFP (MOI 0.2) and viral replication measured at 48hpi using an Incucyte® Live-Cell Imaging System. GFP signal was normalized per image and cell confluence (mean ± SEM, *n* = 4). UF = uninfected.

### Inhibition of RSV-F mediated cell-cell fusion by prolyl-hydroxylase inhibitors

To elucidate the potential mechanism of action of the PHIs on the RSV life cycle, we examined RSV-F driven cell-cell fusion using a split GFP-luciferase reporter cell-cell fusion system, which uses a doxycylcine inducible expression of RSV-F in an effector cell line, combined with a target cell line stably expressing the corresponding half of the reporter (**Fig.3a**) ^43^. Treatment with Daprodustat and Molidustat resulted in a dose-dependent inhibition of RSV-F cell-cell fusion with estimated IC_50_ values of 11 and 42 µM, respectively (**Fig.3b**). To ensure that this activity was RSV-F dependent and not explained by modulation of the split GFP-luciferase reporters, we co-transfected the constructs encoding the half reporters into cells and their activity was not affected by Daprodustat or Molidustat treatment (**Supplementary Fig.7**). To investigate whether the PHIs interfere with RSV-F expression in the effector cells, we measured cell surface RSV-F expression by flow cytometry and treatment had a negligible effect (**Fig.3c**). Furthermore, immunofluorescence analysis indicated that none of the PHIs caused gross changes to the cellular distribution of RSV-F in infected Calu-3 cells (**Fig.3d**). Collectively, these data show that PHIs inhibit RSV-F driven cell-cell fusion.

**Figure 3.**
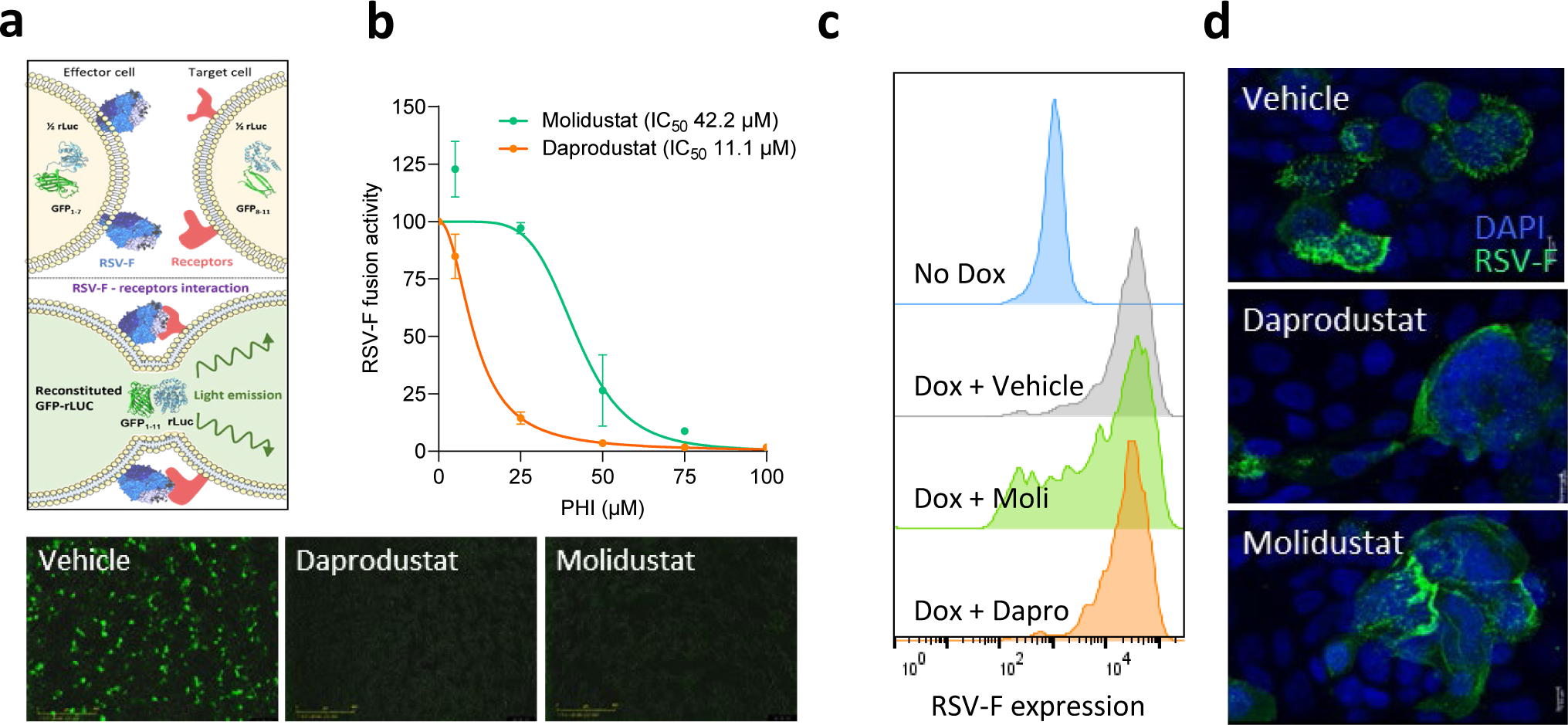
Inhibition of RSV-F mediated cell-cell fusion by prolyl-hydroxylase inhibitors. **a**. HEK293T effector cells (expressing beta strands 1-7 of GFP), engineered for a doxycycline-inducible RSV F protein, were cocultured with HEK293T target cells (expressing beta strands 8-11 of GFP). **b**. The inhibitory activity of Daprodustat or Molidustat was measured as green intensity signal 48h after co-culture (mean ± SEM, n = 3). **c.** HEK293T cells were treated with PHI (50 μM) in the presence of doxycycline and F expression analysed by flow cytometry using an APC-conjugated anti-RSV-F antibody at 24hpi. **d**. Calu-3 cells were infected with RSV (MOI 1) and at 48hpi RSV-F expression imaged.

### Hypoxia and prolyl-hydroxylase inhibitors reduce nucleolin expression via regulating miR-494

We next investigated whether PHIs regulate expression of the reported entry host factors for RSV: ICAM1 ^32^, EGFR ^33^ and NCL ^34^ that may contribute to their anti-viral activity. Analysis of our earlier published transcriptomic data of hypoxic or Roxadustat treated Calu-3 cells ^24^ showed a significant reduction in NCL gene expression (**Fig.4a**). We validated our RNA-seq data and showed that NCL mRNA and protein expression were reduced in Calu-3 cells cultured under low oxygen conditions (1% oxygen) (**Fig.4b**). As a control we confirmed low oxygen induction of NDRG1 gene expression. Daprodustat treatment reduced NCL mRNA and protein expression as assessed by immunoblotting (**Fig.4c**) and flow cytometric analysis of Calu-3 and HEp-2 cells (**Fig.4d, Supplementary Fig.8**). As HIFs are well established transcription activators and generally considered positive regulators of gene expression ^44^, we hypothesised this phenotype of reducing NCL expression is mediated by downstream HIF-signaling and potentially through the regulation of microRNAs. An earlier study reported that NCL transcripts were regulated by microRNA-494 (miR-494) ^45^. Notably, miR-494 has a HRE in its promoter (**Supplementary Fig.9**) and was previously reported to be induced under hypoxic conditions ^46,47^. To examine this hypothesis, we measured miR-494 levels in Calu-3 cells cultured at 1% oxygen or treated with Daprodustat and found a robust induction (**Fig.4e**). Moreover, treatment with a miR-494 mimic reduced NCL protein expression and significantly reduced RSV infectivity (**Fig.4f**), supporting HIF regulation of NCL expression through induction of miR-494.

**Figure 4.**
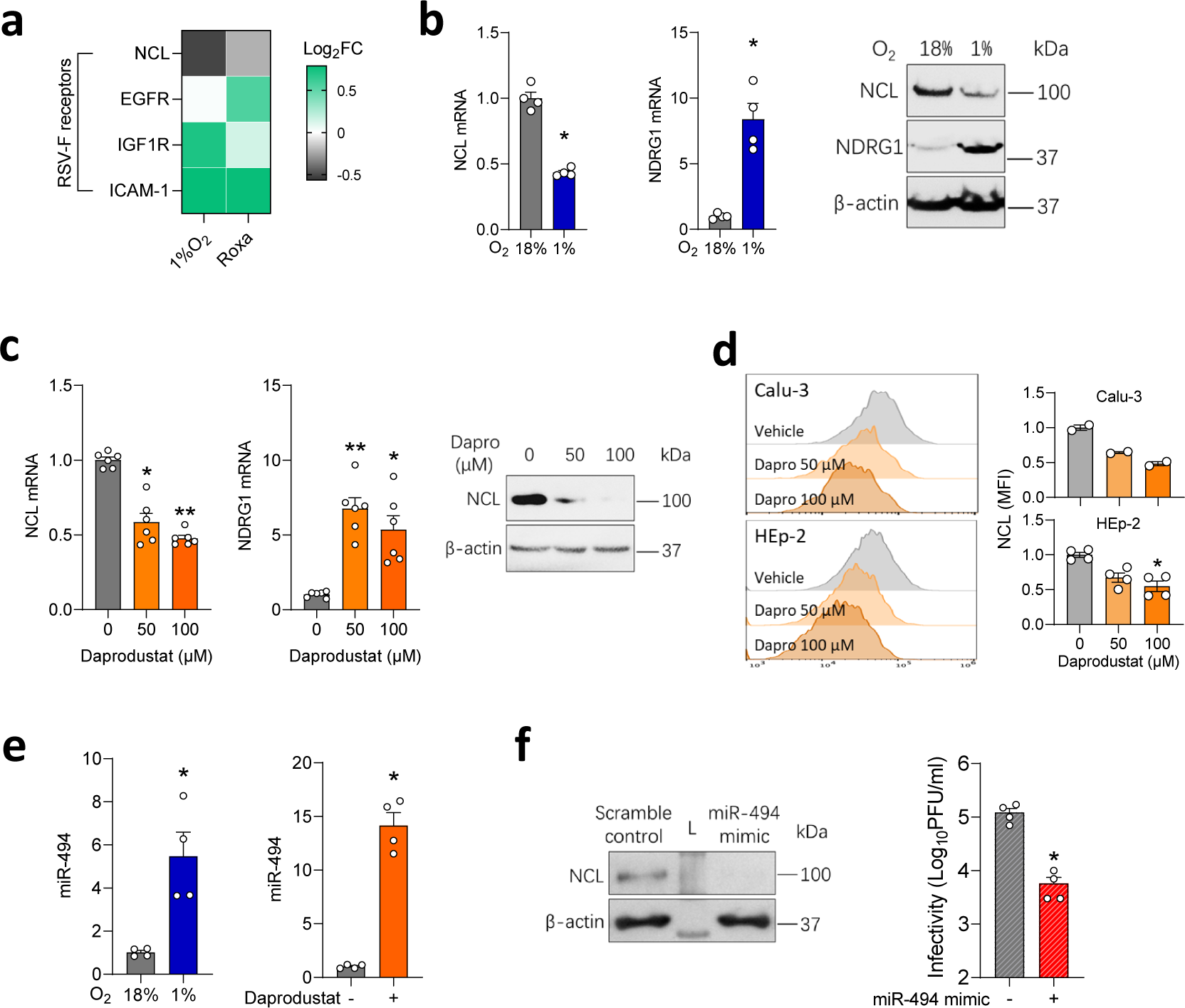
Hypoxia or prolyl-hydroxylase inhibitors inhibit nucleolin via regulating miR-494. **a**. Calu-3 cells were cultured at 18% or 1%O_2_ or treated with Roxadustat (50 µM) for 48h and cellular RNA sequenced. Differential expression analysis was performed to interrogate the expression of host entry factors that regulate RSV fusion, where the heat-map denotes Log_2_ Fold change (FC). **b**. Calu-3 cells were cultured at 18% or 1% O_2_ for 48h and NCL and NDRG1 mRNA or protein expression measured by qPCR or immunoblotting, respectively (mean ± SEM, *n* = 4, Mann–Whitney test, Two-sided). **c**. Calu-3 cells were treated with Daprodustat for 48h and NCL mRNA and protein expression determined by qPCR and immunoblotting, respectively (mean ± SEM, *n* = 6, One-way ANOVA with multiple comparisons, Two-sided). **d**. Calu-3 or HEp-2 cells were treated with Daprodustat for 48h and total NCL protein expression determined by flow cytometry (mean ± SEM, n = 2-4, One-way ANOVA with multiple comparisons, Two-sided). **e**. Calu-3 cells were cultured at 18% or 1% O_2_ or treated with Daprodustat (50 µM) for 48h and miR-494 expression measured by qPCR (mean ± SEM, *n* = 4, Mann–Whitney test, Two-sided). **f.** Calu-3 cells were pre-treated with a miR-494 mimic (1 µM) for 24h before infecting with RSV (MOI 0.2) and intracellular infectivity determined at 48 hpi (mean ± SEM, *n* = 4, Mann–Whitney test, Two-sided). L = Molecular weight ladder.

### Daprodustat limits RSV infection and pulmonary inflammatory responses in mice

To assess the *in vivo* activity of Daprodustat, we administered the drug twice daily at a dose of 10 mg/kg or 30 mg/kg to mice and euthanised the animals after 4 days of treatment. This regimen was based on previous dosing protocols in mice ^23,48^. We confirmed the drug was well-tolerated as there was no evidence of weight loss or histopathological changes in the lungs (**Fig.5a; Supplementary Fig.10**). As HIF expression following systemic PHI treatment is transient and difficult to detect ^49^, we evaluated Daprodustat efficacy by assessing HIF activation of erythropoietin stimulated erythrocytosis by measuring immature red blood cells (reticulocytes). Blood smears from terminal blood samples showed increased reticulocyte counts compared to vehicle, consistent with effective drug treatment (**Fig.5b**). To establish whether oral delivery of Daprodustat was effective in the lower respiratory tract we showed activation of endothelin 1 (*Edn1*) a gene reported to be HIF regulated in the respiratory tract^50^ (**Fig.5b**). In addition we observed reduced NCL gene expression in the lung of Daprodustat treated mice (**Fig.5b**).

**Figure 5.**
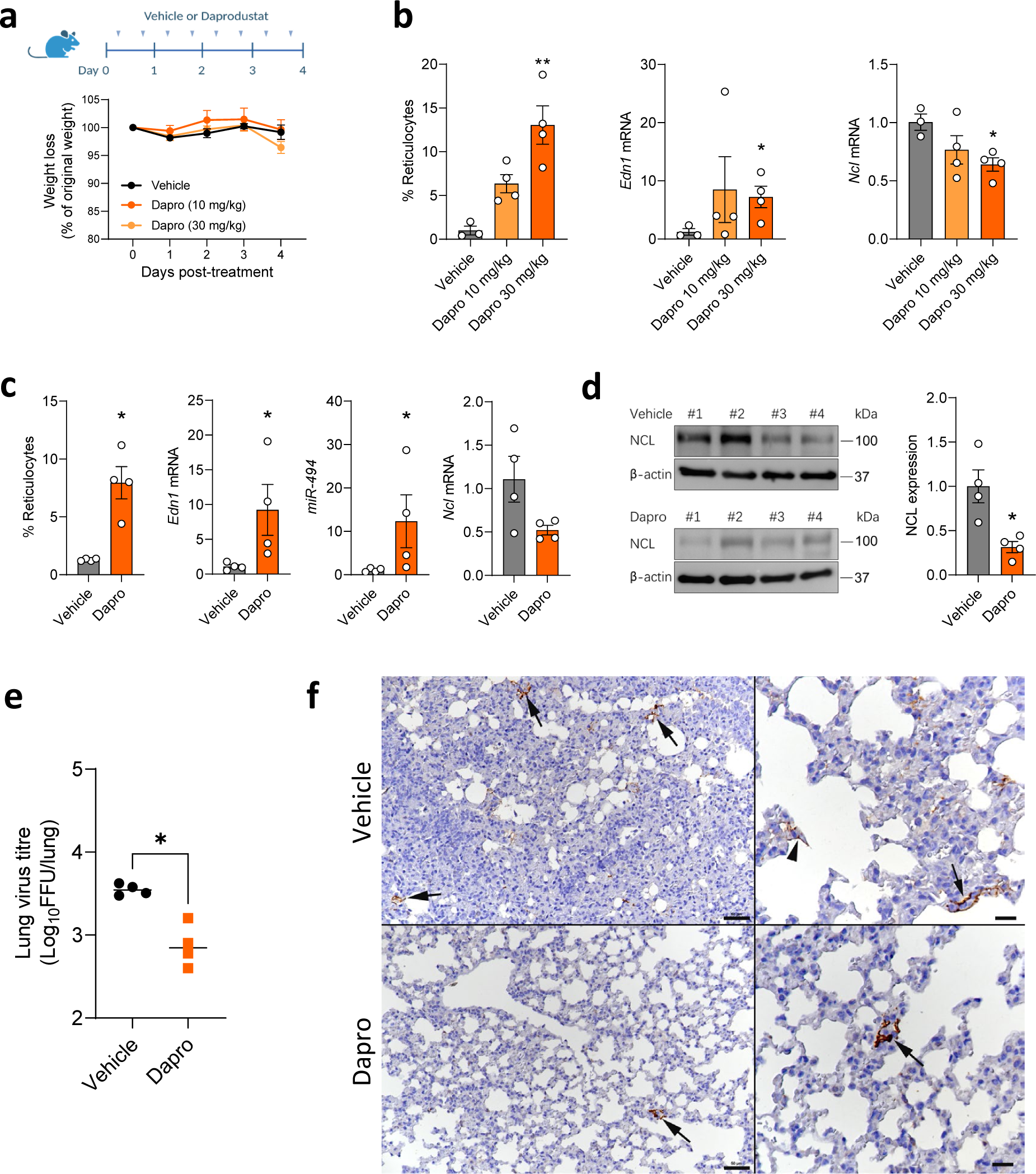
Daprodustat treatment reduces RSV infection in mice. **a**. BALB/c mice (6-8 weeks old, female) were treated with vehicle (1% methylcellulose) or Daprodustat (10 or 30 mg/kg), twice daily via oral gavage for 4 days. **b**. Reticulocyte counts were measured and RNA extracted from the lung and *Edn1* and *Ncl* mRNA levels measured by qPCR (mean ± SEM, vehicle n = 3; Daprodustat n = 4, One-way ANOVA with multiple comparisons, Two-sided). **c**. BALB/c mice (6-8 weeks old, female) were treated with vehicle (1% methylcellulose) or Daprodustat (30 mg/kg), BID, via oral gavage 24h prior to intranasal infection with RSV A2 (1×10^7^ PFU) and treated with drug as stated in (a) for 4 days. At the end of the experiment, reticulocyte counts were performed, and RNA extracted from the lung for qPCR measurement of *Edn1*, *miR-494* and *Ncl* mRNA levels (mean ± SEM, *n* = 4, Mann–Whitney test, Two-sided). **d**. Lung homogenates were assessed for NCL protein expression by immunoblotting and densitometric analysis quantified. NCL in individual samples were normalised to their β-actin loading controls (mean ± SEM, *n* = 4, Mann–Whitney test, Two-sided). **e**. RSV infectivity in lung homogenate was measured by a focus forming assay (mean ± SEM, n = 4, Mann–Whitney test, Two-sided). **f.** Representative images of lungs from mice at 4 dpi. The lung of a vehicle treated mouse exhibits several small patches of alveoli with RSV antigen positive pneumocytes (left: arrows). A higher magnification (right) shows infected type I pneumocytes (arrowhead) and type II pneumocytes (arrow). The lung of a Daprodustat (30 mg/kg) treated mouse shows a single patch of viral antigen expressing pneumocytes (left: arrow). The higher magnification of the patch (right; arrow) confirms that both type I and II pneumocytes are infected. Immunohistology, hematoxylin counterstain. Bars = 50 µm (left) and 20 µm (right).

To evaluate the anti-viral activity of Daprodustat, we infected mice intranasally with RSV and treated with the drug at 30 mg/kg using the same dosing schedule as above, followed by euthanasia at 4 dpi. We observed a minimal weight loss in the vehicle or Daprodustat treated RSV infected groups. We confirmed HIF pathway activation by measuring reticulocyte counts and increased *Edn1* expression in the Daprodustat treated group. We observed a reduction in *Ncl* mRNA and protein along with an increase in *miR-494* levels in the drug treated group compared to the vehicle controls (**Fig.5c-d**). In Daprodustat treated there was a significant reduction in the infectious RSV titer in the lung (**Fig.5e**). Immunohistological staining of the lungs for RSV antigen detected virus infected type I and II pneumocytes in low numbers of random disseminated alveoli, and with lower frequency in the drug treated mice (**Fig.5f**).

RSV infection was accompanied by a generally mild inflammatory response, represented by mononuclear perivascular and/or peribronchial infiltrates (**Fig.6**). This was was consistently more extensive in the vehicle treated mice where it was accompanied by leukocyte rolling, subendothelial accumulation and infiltration of vascular walls (vasculitis), with circular perivascular accumulation in several layers. In Daprodustat treated mice, there was only minimal leukocyte recruitment, with often fragmentary and thin perivascular accumulation. The dominant cells recruited from the vessels were monocytes (Iba1+) and T cells (CD3+), intermingled with rare neutrophils (Ly6G+). In addition, all lungs exhibited rare to a few small focal parenchymal areas with activated type II pneumocytes and a few infiltrating macrophages, lymphocytes and neutrophils (**Fig.6**).

**Figure 6.**
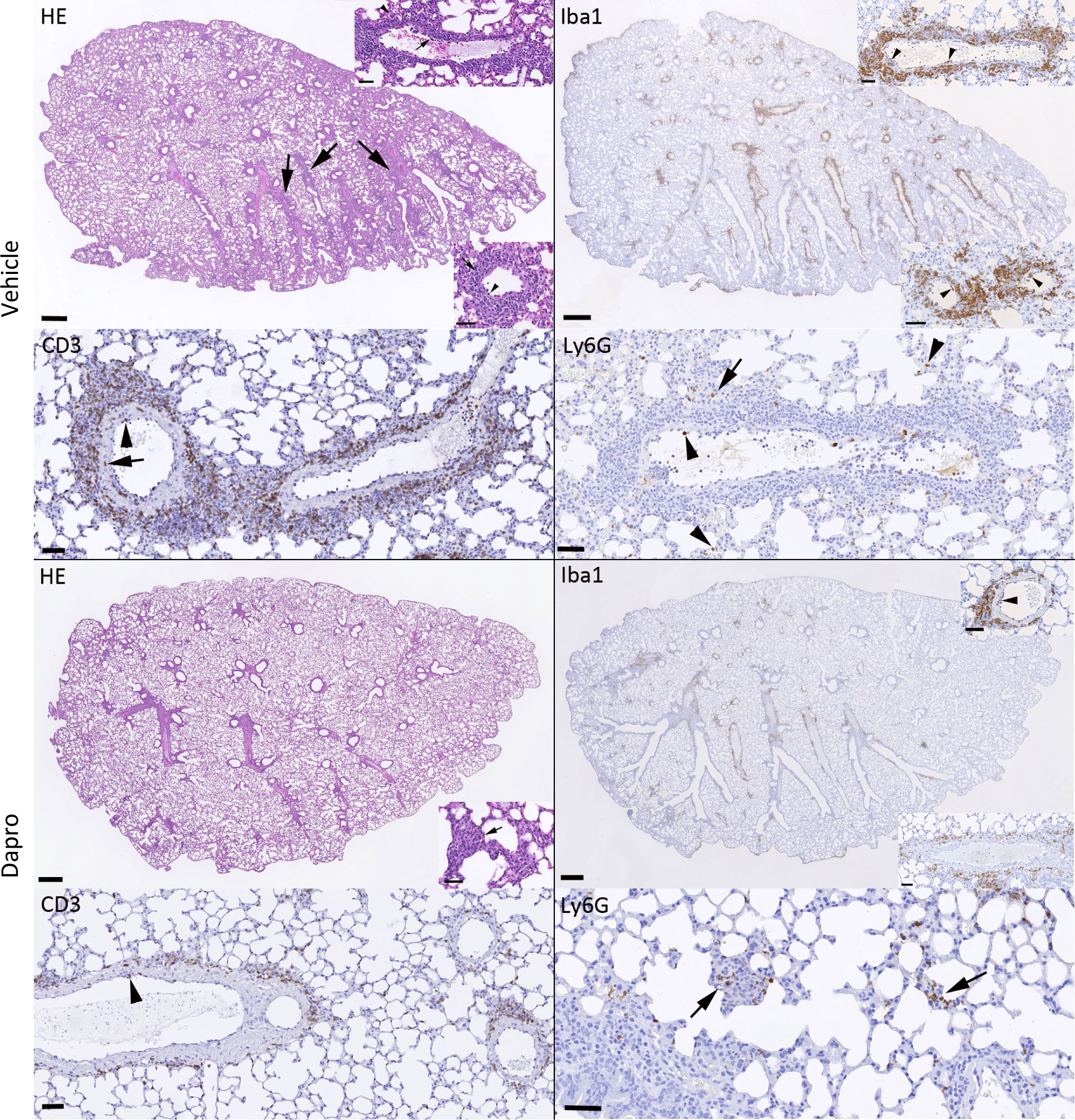
Daprodustat treatment reduces the RSV-associated pulmonary inflammatory responses in mice. Representative images of lungs from mice at 4 days post RSV infection. As shown in the HE stained section, the lung of a vehicle treated mouse exhibits mild perivascular and peribronchial mononuclear infiltrates (arrows) associated with leukocyte rolling and emigration (bottom inset, arrowhead) and subendothelial leukocyte aggregates in particular in muscular veins (top inset, arrowhead) consistent with vasculitis. The infiltrates are dominated by monocytes/macrophages (Iba1+) which are also found rolling and attached to the endothelium (bottom inset: arrowheads) and within the subendothelial infiltrates (top inset: arrowheads). T cells (CD3+) are numerous and present rolling (arrowhead) and within the vascular wall (arrow). There are a few neutrophils (Ly6G), within the lumen of vessels and capillaries (arrowheads) and occasionally within the small parenchymal infiltrates (arrow). After Daprodustat treatment, the inflammatory response is limited to very mild perivascular mononuclear infiltrates and rare small focal parenchymal infiltrates (HE stain: inset, arrow). The infiltrate shows a similar composition as in the vehicle treated mice. It is dominated by monocytes/macrophages (Iba1+) that are occasionally found also in vascular walls (top inset: arrowhead), followed by T cells (CD3+) that are occasionally seen rolling along the vascular endothelium (arrowhead). Neutrophils (Ly6G+) are mainly seen in the rare small parenchymal infiltrates (arrows). Bars = 500 µm (whole section images) and 50 µm (all other images).

RNA-sequencing of the lung from vehicle or Daprodustat treated mice confirmed an activation of hypoxic-regulated pathways including glycolysis and showed evidence of increased interferon (alpha/gamma), TNF and TGFbeta signalling, consistent with an increase in immune signalling pathways (**Supplementary Fig.11**). Cholesterol is an essential lipid in the life cycle of virtually all viruses ^51^, including RSV ^52^ and it is noteable that Daprodustat suppressed cholesterol homeostasis pathways (**Supplementary Fig.11**). These transcriptomic analyses support a wide-ranging role for Daprodustat to perturb immune signalling pathways that will impact RSV replication and are worthy of futher study. Together these data demonstrate that Daprodustat treatment reduced the frequency of RSV infected pneumocytes and infectious viral burden in treated mice.

## DISCUSSION

The interplay between HIF-signalling pathways and human pathogens has been studied for multiple viruses ^53^ and more recently in the context of respiratory viral infections ^23,24,54^. RSV, Influenza and SARS-CoV-2 replicate within the respiratory tract, leading to cell damage and triggering an inflammatory response that disrupts oxygen distribution and contributes to local hypoxia. Influenza A virus infection of mice with HIF-1α knock-out in type II alveolar epithelial cells ^55^ showed a significant increase in viral replication, highlighting an antiviral role for HIF-1α in this respiratory disease. The pathogen specific mechanisms and interactions with HIFs may vary depending on the virus, and understanding this relationship will underpin the development of effective treatments and interventions for respiratory viral infections.

Hypoxic conditions or treatment with PHIs to stabilise HIFs suppressed RSV replication. We confirmed this antiviral phenotype in Calu-3 and HEp-2 epithelial cell lines, in BEAS-2B that are derived from normal human bronchial epithelium and ALI PBEC cultures. RSV can induce robust syncytia formation *in vitro* and our real-time imaging showed that PHIs inhibit cell-cell fusion and spread of a GFP-expressing recombinant RSV. Moreover, the addition of Palivizumab further enhanced this inhibition. Through an RSV F protein-driven cell-cell fusion assay, we established that viral spread was a primary target of the PHIs (Daprodustat or Molidustat), and identified the downregulation of the RSV receptor nucleolin as the likely mechanism. We demonstrated the sensitivity of RSV to Daprodustat-mediated nucleolin knockdown *in vivo* using an experimental mouse model of RSV infection, and showed upregulation of the HIF-regulated microRNA - miR-494. Overall, these findings provide a mechanistic link between HIF-mediated transcription and inhibition of RSV infection (**Fig.7**). PHIs were developed as erythropoiesis-stimulating agents for treating anaemic patients with chronic kidney disease. One limitation with repurposing this class of drugs is their primary pharmacological activity to increase hemoglobin concentrations^56^. However, there are several PHIs available with different selectivities^57^ and evaluating their antiviral activity in an RSV mouse model would be worth investigating.

**Figure 7.**
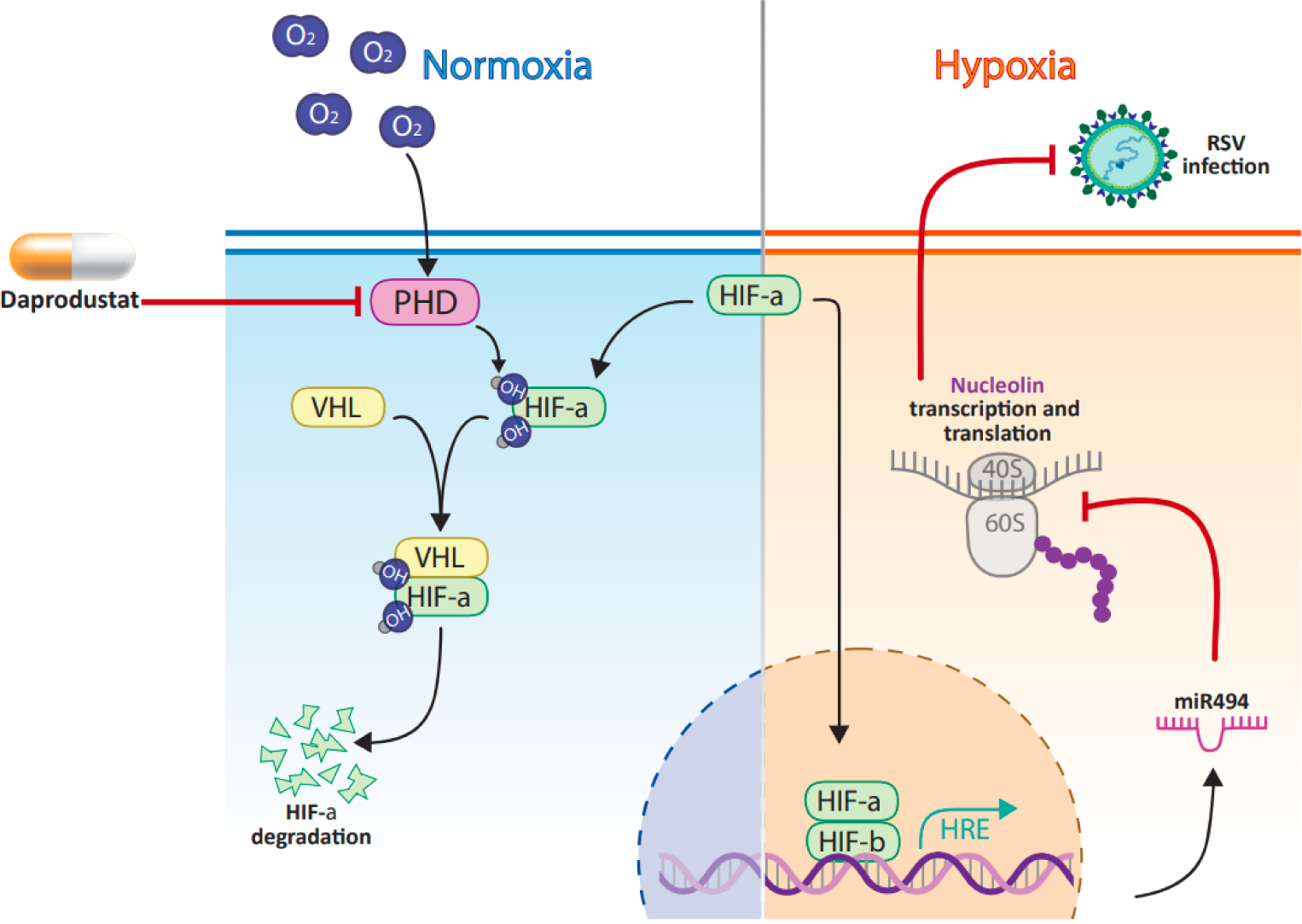
Model of HIF signalling regulation of RSV infection. Hypoxia or Daprodustat treatment induces HIF expression and inhibits RSV infection by upregulation of miR-494, leading to a reduction of the viral entry receptor nucleolin.

Earlier studies reported that RSV infection induced glycolysis via activating HIF-1α expression and the HIF-1α inhibitor PX-478 reduced virus replication^36,37,58^. PX-478 has been reported to reduce HIF-1α mRNA levels and translation under both normoxic and hypoxic conditions^59^, which may complicate the interpretation of these studies in assessing the effect(s) of HIF-signalling on the RSV life cycle. We did not observe HIF-1α protein or activity in RSV infected Calu-3 cells and noted limited evidence of HIF regulated gene expression in the infected ALI-PBEC cultures. These differing results may reflect the variable proliferative and metabolic status of the infected target cells that can influence their glycolytic rate and endogenous HIF-signallling pathways. Recent reports demonstrate that RSV encoded matrix protein induces a perinuclear clustering of the mitochondria and infection perturbs mitochondrial function, resulting in increased reactive oxygen species (ROS) ^60,61^. The impact of ROS on the transcriptional and translational regulation of HIF-1α are complex and can activate different genes to those seen under hypoxia (reviewed in ^62^). Hypoxic conditions have been reported to induce perinuclear clustering of mitochondria to create an oxidant-rich environment that is important for downstream gene regulation and HIF-signaling ^63–66^. Further investigations are warranted to better understand the interplay between RSV induced changes in cellular metabolism and HIF signalling pathways.

We demonstrate that miR-494 is induced under hypoxia and Daprodustat treatment. Moreover, we found that a miR-494 mimic reduced NCL expression and inhibited RSV infection, revealing a novel role of oxygen-sensing microRNAs in regulating RSV infection. However, additional factors may influence NCL expression. The interaction between HIF and the transcriptional regulator c-myc may also suppress NCL expression. Myc regulates the transcription of NCL via binding e-boxes in the gene promoter ^67^, however Myc requires MAX, its binding partner, to activate gene expression ^68^. In hypoxic conditions, MAX is sequestered by HIF-1α, which impairs its ability to transcribe target genes including NCL. RNA-seq analysis of the lung from vehicle or Daprodustat treated mice showed an enrichment in Myc target pathways (**Supplementary Fig.11**). It is likely that other oxygen sensitive pathways, such as chromatin conformation and DNA methylation may further modulate NCL expression in hypoxic conditions. NCL was reported to interact with Influenza virus non-structural protein 1 and to regulate late viral gene expression ^69,70^, and to also mediate SARS-CoV-2 replication ^71^, suggesting a wide-ranging impact for the hypoxic regulation of NCL in respiratory infections.

Tayyari et al identified NCL as a human RSV receptor using several experimental approaches including siRNA knockdown, antibody interference, overexpression in non-permissive cells and *in vivo* challenge studies ^34^. We confirmed a role for NCL in RSV infection of Calu-3 cells by showing reduced infection following transient siRNA mediated knock-down of NCL expression (**Supplementary Fig.12**). NCL ligands such as pseudopeptide HB-199, midkine or pleiotrophin have been proposed as potential therapeutics against RSV ^72–74^. Mastrangelo *et al* reported that the NCL binding anti-cancer drug, AS1411, could ameliorate RSV infection ^75^. More recently, the same authors reported that RSV-F binds the RNA binding domains 1 and 2 of NCL and peptides from this region could inhibit RSV infection ^76^. As NCL has been reported to interact with other viruses including parainfluenza type 3 ^77^, enterovirus 71 ^78^, Crimean–Congo haemorrhagic fever virus ^79^ and adeno-associated virus type 2^80^, our observation showing HIF/miR494 regulation of NCL may be widely applicable to many viruses.

NCL is often over-expressed in cancer cells and although the mechanisms for its cell surface expression are not well characterised, NCL is thought to cluster at the membrane through cytoskeletal interactions ^81^. Deregulation and over-expression of NCL on the cell surface have been associated with lower survival rates, potentially defining it as a biomarker. Moreover, the ability of NCL to internalise a range of ligands in a calcium-dependent manner has made it an attractive target for therapeutic intervention (reviewed in ^82^). It is intriguing to speculate that hypoxic conditions or pharmacological stabilisation of HIFs, as modeled in our study, may disrupt cancer-specific surface NCL overexpression. Nucleolin is widely targeted for intracellular drug delivery, and this newly identified mechanism for its downregulation may shed further light on the role of this protein in cancer development, particularly considering the hypoxic nature of many solid tumors. It is worth noting that while the majority of our work was performed in Calu-3 cells, a non-small-cell lung cancer line, we observed similar phenotypes in BEAS-2B cells and in ALI-PBEC which are derived from normal bronchial epithelium.

Using an established RSV mouse model we show that Daprodustat treatment reduced the level of infectious virus and this was accompanied by a reduced frequency of RSV antigen expressing pneumocytes. The frequency of infected cells in the lung at 4dpi was low and this may reflect the semi-permissive nature of hRSV infection of mice (reviewed in^83^). Despite the low frequency of infection, RSV induced a similar inflammatory response as seen with other respiratory viruses, such as SARS-CoV-2 and IAV, with clear evidence of leukocyte recruitment and monocyte and T cell driven vasculitis and perivascular/peribronchial infiltration^84^. Daprodustat treatment dampened these inflammatory responses, although the recruitment of neutrophils was maintained, in line with earlier reports on hypoxia driving neutrophilic inflammation^85^. Preliminary experiments in mice infected with the genetically related pneumovirus of mice (PVM), a natural pathogen of rodents, support our findings with hRSV and showed a reduced frequency of infected cells and a less intense inflammatory response following Daprodustat treatment. Further studies are warranted to identify the inflammatory cells and their localisation with RSV infected epithelial cells during the acute and resolution phases of disease. Reassuringly, these observations align with our recent report that treating hamsters with the PHI Roxadustat reduced SARS-CoV-2 infection and epithelial damage ^24^, highlighting a conserved role for HIFs to suppress respiratory viruses.

In summary, our study has uncovered a novel mechanism for HIF to downregulate NCL, the cell surface receptor for RSV, that is mediated through the transcriptional activation of miR-494. Whilst our findings support a role for HIFs to limit RSV RNA replication understanding the detailed molecular mechanisms is beyond the remit of the current study. A recent report that hypoxia promotes JMJD6 mediated hydroxylation of proteins in regions predicted to be unstructured and associated with membraneless organelle formation^86^ provides a potential mechanism for HIF-independent hypoxic pathways to perturb RSV RNA replication and is worthy of investigation. This study highlights a role for HIF-signalling to limit multiple aspects of RSV infection and associated inflammatory responses.

## MATERIALS AND METHODS

### Cells and reagents

All cells, including co-cultures, were cultured at 37°C and 5% CO_2_ in a standard culture incubator and exposed to hypoxia using an atmosphere-regulated workstation set to 37°C, 5% CO_2_ and 1% O_2_ (Invivo 400, Baker-Ruskinn Technologies). Calu-3 cells were cultured in Advanced DMEM (Sigma-Aldrich) supplemented with 10% fetal bovine serum (FBS), 2mM L-glutamine, 100 U/mL penicillin and 10 mg/mL streptomycin (Invitrogen). HEp-2 and BEAS-2B cells were cultured in DMEM (Sigma-Aldrich) with the same supplements as above. Human PBECs were obtained using flexible fibreoptic bronchoscopy under light sedation with fentanyl and midazolam from healthy control volunteers and all participants provided written informed consent. The study was reviewed by the Oxford Research Ethics Committee B (18/SC/0361). Airway epithelial cells were cultured in Airway Epithelial Cell medium (PromoCell, Heidelberg, Germany) in submerged culture. PBECs were cultured on PureCol-coated 0.4 μm pore polyester membrane permeable inserts (Corning) in serum-free airway epithelial cell media, brought to air-liquid interface, replacing basal media with ALI media (StemCell Pneumacult). The media was exchanged every 2 days and apical surfaces washed with PBS weekly to disperse accumulated mucus for a minimum of 6 weeks.

Real-time luciferase activity was acquired at 30 min intervals using a ClarioStar plate reader (BMG). HEK293T cells stably expressing Lenti-rLuc-GFP 1–7 and RSV-F protein, or separately, Lenti-rLuc-GFP 8–11, were used for fusion assays and maintained using DMEM (Sigma-Aldrich) supplemented with 10 % FBS (Life Science Production), 1 % NaP (Sigma-Aldrich) and 1 % Pen-Strep (10000 U ml−1; Life Technologies Ltd). Daprodustat (GSK: 1278863), Molidustat (Bay 85-3934), Roxadustat (FG-4592) were obtained from MedChemExpress. For IH the following antibodies were used: goat anti-RSV (AB1128; Millipore), goat-HRP (ab6741; Abcam) and Envision+System HRP Rabbit (K4003; Agilent Dako).

### siRNA silencing

Silencing RNA for NCL (human NCL silencer select) and siRNA controls were obtained from Thermo Fisher. For siRNA-mediated silencing, NCL siRNA or a scrambled control at 25 nM were transfected using DharmaFECT 4 (Thermo Fisher). 24h post transfection, Calu-3 were infected with RSV-GFP and monitored using Incucyte S3 live cell imaging. 48h post transfection cells were harvested for western blotting.

### Cell-cell fusion assay

HEK293T Lenti rLuc-GFP 1–7 engineered to express RSV-F (effector cells) and HEK293T Lenti rLuc-GFP 8–11 (target cells) were co-cultured in a 96-well plate in the presence of doxycycline (1 µg/mL), to induce the expression of RSV-F by effector cells ^87^. Each cell line was diluted to dispense 2×10^4^ cells/well in a final volume of 100 μL and PHIs were added to the culture media. After 48h, GFP signal was quantified using IncuCyte S3 live cell imaging system (Essen BioScience).

### RSV infection and quantification of virus infectivity

Human RSV subtype A (A2 strain) was grown in HEp-2 cells and cell lysates were concentrated using Vivaspin® 20, 10,000 MWCO PES columns (Sartorius). In brief, cells were infected at MOI 0.2 for 2h, the inoculum was removed, cells washed with PBS and cultured in DMEM 2% containing 5% FBS. After 5 days, cells were harvested and cellular lysates clarified and concentrated using Vivaspin® 20. The aliquoted viral stock was snap frozen in liquid nitrogen and stored at -80°C.

Calu-3, HEp-2 or BEAS-2B cells were infected with RSV at an MOI of 0.2 for 2h, the viral inocula removed by washing the cells three times in PBS and the cultures maintained in growth media. PBEC-ALI cultures were washed with PBS and infected apically with RSV-A2 (an estimated MOI of 5) for 2h at 37°C. Daprodustat was added in the basal medium and shed virus harvested at 48hpi by washing the cultures with 50 µl of medium followed by a standard focus forming assay using HEp-2 cells. For viral kinetics, Calu-3 were seeded at 5×10^4^ cell/well in a clear flat-bottomed 96 well-plate. Cells were allowed to grow to 80% confluence, and infected with RSV-GFP. After removal of the inoculum, plates were imaged every 8h using an IncuCyte S3 live cell imaging system. GFP expression was determined using the total integrated intensity metric function in the IncuCyte S3 software.

Samples from infected cells and lung homogenates from infected mice were serially diluted 1:10 and used to inoculate monolayers of HEp-2 cells for 2h. Inocula were removed and replaced with DMEM containing 1% FCS with a semi-solid overlay consisting of 1.5% carboxymethyl cellulose (Sigma-Aldrich). Cells were incubated for 4-5 days, after which cells were fixed in 4% PFA, stained with 0.2% crystal violet (w/v) and visible plaques enumerated, where the limit of detection of this assay was 10 focus forming units (FFU)/mL.

### RSV minigenome

Calu-3 cells were seeded in 24-well plates, inoculated with Fowlpox-T7 and the following day transfected with plasmids encoding the RSV replicase: phosphoprotein (P); nucleoprotein (N); polymerase (L); anti-transcription-termination factor (M2-1) and a subgenomic human RSV A2 luciferase replicon^38^. Transfected cells were cultured at 37°C for 24 h, treated with drugs and 48h later lysed and luciferase determined as a measure of hRSV replication and transcription.

### qPCR quantification

Viral RNA was extracted from infected cells using a RNeasy kit (Qiagen) according to the manufacturer’s instructions. Lung tissue from infected mice were homogenised in Trizol and extracted according to the manufacturer’s instructions. Viral or cellular RNA was determined by qPCR using a Roche Light Cycler 96.

### Primers

Oligonucleotide sequences (Life Technologies)

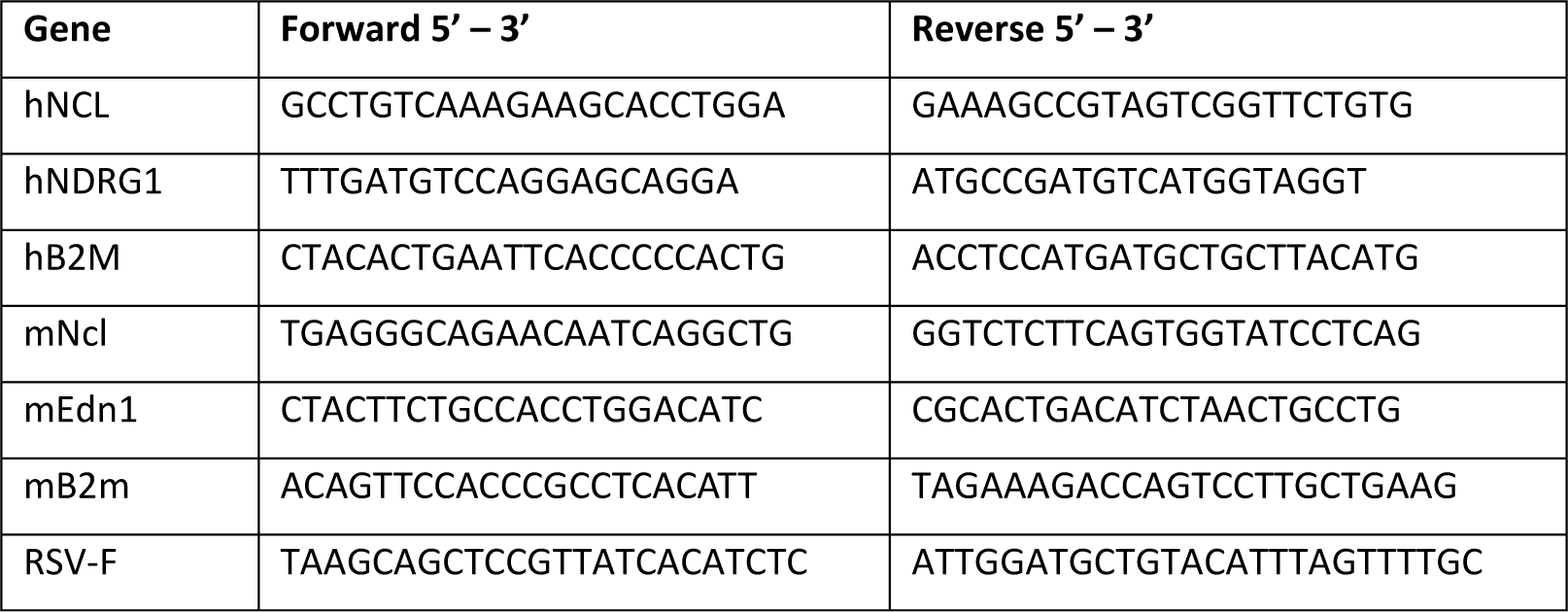

### Immunoblotting

Cell lysates were prepared by washing cells with phosphate buffered saline (PBS), then lysed in Igepal lysis buffer (10mM Tris pH 7.5, 0.25M NaCl, 0.5% Igepal) supplemented with Complete TM protease inhibitor cocktail (Roche) at 4°C for 5 min, followed by clarification by centrifugation (3 min, 12,000 rpm). Supernatant was mixed with Laemmli sample buffer and boiled at 100°C, separated by SDS-PAGE and proteins transferred to polyvinylidene difluoride membrane (Immobilon-P, Millipore). Membranes were blocked in 5% milk in PBS/ 0.1% Tween-20, incubated with anti-HIF-1α (BD Transduction Laboratories, clone#610959), anti-NCL (abcam, clone# 22758), anti-NDRG1(Cell signaling, clone#5196) or anti-β-actin (Sigma, clone#A5441) primary antibodies and appropriate HRP-conjugated secondary Mouse antibodies (DAKO, clone#P0447) and Rabbit antibodies (Cytiva, clone#NA934V). Chemiluminescence substrate (West Dura, 34076, Thermo Fisher Scientific) was used to visualize proteins using a ChemiDoc XRS+ imaging system (BioRad). Densitometric analysis was performed using ImageJ software (NIH).

### Flow cytometry

HEK293T effector cells were plated in a 12 well-plate in presence of PHI and doxycycline. The following day, they were stained with an APC conjugated anti-RSV-F antibody and with LIVE/Dead™ Fixable Aqua (Life Technologies) for 30 min at 4°C. Cells were analysed on a MACSQuant Analyzer flow cytometer (Miltenyi Biotec) and data analyzed using FlowJo (TreeStar). HEp-2 or Calu-3 cells were seeded in a 6 well-plate and incubated with Daprodustat (50 mM or 100 mM) for 48h. Cells were trypsinised, stained with LIVE/Dead™ Fixable Aqua (Life Technologies), permeabilized for 15min at room temperature (0.1% Triton X-100 in PBS), fixed with 4% paraformaldehyde (PFA, Sigma) for 10min at room temperature and blocked for 20min on ice (0.5% BSA, 0.5% tween in PBS). Cells were stained in darkness with primary nucleolin antibody (abcam, ab129200) for 20 min on ice, followed by staining with in fluorescent dye conjugated secondary antibody (goat anti-rabbit Alexa 488nm, Thermo Fisher, A11034) for 20min on ice. Data was acquired using an Attune NxT flow cytometer (Thermo Fisher) and analyzed using FlowJo (TreeStar).

### Confocal immunofluorescence microscopy

Infected cells were fixed with 4% paraformaldehyde (PFA; Sigma) in PBS for 15 min, blocked with 20mM glycine in PBS and permeabilized with 0.5% Triton X-100 in PBS for 5 min. Cells were incubated with anti-RSV-F primary antibody for 1h at room temperature. After incubation, they were washed and incubated with Alexa Fluor secondary antibodies (Life Technologies) for 1h at room temperature. After washing, slides were mounted with Fluoromount G (SouthernBiotech) containing 4′,6-diamidino-2-phenylindole (DAPI) for nuclei staining. Cells were imaged on a Leica TCS SP5 confocal microscope using the 488-nm laser line for the appropriate dyes and a 63× oil immersion objective.

### Animals and Daprodustat treatment

Animal work was approved by the local University of Liverpool Animal Welfare and Ethical Review Body and performed under UK Home Office Project Licence PP4715265. Female BALB/c mice (BALB/cAnNCrl), 8-10 weeks of age were purchased from Charles River and maintained under SPF barrier conditions in individually ventilated cages. Mice were divided in two groups for treatment with vehicle or Daprodustat (n = 4-6 per group). Animals were treated with 10mg/kg or 30mg/kg of Daprodustat (MedChem Express) by oral gavage. Drug was dissolved in 99% double distilled H_2_O, 0.5% methyl cellulose and 0.5% Tween-80, administered twice daily. Animals were euthanised on day 4 post-treatment and tissues collected for histological examination. Reticulocyte counts were quantified by staining terminal blood samples with 0.1% Brilliant Cresyl Blue. Animal work was approved by the local University of Liverpool Animal Welfare and Ethical Review Body and performed under UK Home Office Project Licence PP4715265.

### RSV-Daprodustat challenge experiments

Two identical experiments were performed. For both, female BALB/c mice (8-10 weeks old) were randomly assigned into cohorts (4 or 6 animals). Daprodustat (30mg/kg)/vehicle treatment commenced 24h prior to infection and was maintained until termination of the study 4 days post infection. For infection, mice were anaesthetized lightly with KETASET i.m. and inoculated with 1×10^7^ PFU of RSV-A2 strain. Viral inocula were made in sterile PBS and delivered via intranasal instillation (50 μL total per mouse). Animal weights were monitored daily and visual inspection of all animals carried out twice daily. Animals were euthanised at day 4 post infection and lungs dissected at necropsy for histological and immunohistological examination (left lungs) and virology assays (right lungs). Reticulocyte counts were quantified by staining terminal blood samples with 0.1% Brilliant Cresyl Blue.

### Histological and immunohistological examination

The left lungs were fixed in 10% buffered formalin for 48h, stored in 70% ethanol until processing. Lungs were trimmed (longitudinal section) and routinely embedded in paraffin wax. Consecutive sections (3 µm) were prepared and stained with haematoxylin eosin (HE) for histological assessment and subjected to immunohistology (IH) for the detection of RSV antigen. IH was performed using the horseradish peroxidase (HRP) method. Briefly, after deparaffination, sections underwent antigen retrieval in Tris/EDTA buffer (pH 9) for 20 min at 98°C, followed by incubation with goat anti-RSV (Millipore) diluted in dilution buffer (Agilent Dako) overnight at 4°C. This was followed by blocking of endogenous peroxidase (peroxidase block, Agilent Dako) for 10 min at room temperature (RT) and incubation with goat HRP (Abcam) for 30 min at RT, all in an autostainer (Agilent Dako). Sections were counterstained with haematoxylin. A formalin fixed and paraffin embedded RSV infected (MOI 0.2, 48 hpi) Calu-3 pellet served as positive control; tissue sections incubated without the primary antibody served as negative controls. Further sections were stained for Iba1 (monocytes/macrophage marker), CD3 (T cell marker) and Ly6G (neutrophil marker) following previously published protocols^84,88^.

### RNA-sequencing analysis

Total cellular RNAs were prepared as described above and sequenced using 300bp paired end Illumina sequencing (Novogene, UK). Host reads were mapped to the human transcriptome and differential expression analysis performed using DESeq2. Pathway analyses were performed using Gene Set Enrichment Analysis (GSEA 4.1.0), and the Hallmarks gene sets from the Molecular Signatures Database.

### Quantification and statistical analysis

All data are presented as mean values ± SEM. P values were determined using a Mann-Whitney test (two group comparisons) or with a Kruskal–Wallis ANOVA (multi group comparisons) using PRISM version 8. In the figures * denotes p < 0.05, ** < 0.01, *** <0.001 and **** <0.0001.

## ACKNOWLEDGEMENTS.

The authors are grateful to the laboratory technicians, Histology Laboratory, Institute of Veterinary Pathology, Vetsuisse Faculty, University of Zurich, for excellent technical support. We acknowledge the support of Geraldine Taylor (The Pirbright Institute), Ursula Buchholz (NIAID, NIH) and Karl-Klaus Conzelmann (Max-von-Pettenkofer Institut) for providing recombinant viruses, Tim Hinks (Oxford University, UK) for human PBECs, Julian Hiscox (University of Liverpool) for sharing RSV minigenome reagents and Nadina Wand for critical reading of the manuscript.

## AUTHOR CONTRIBUTIONS

XZ, DB and JAM conceived the project; XZ, GG, PS, AM, HB, JMH, ST, PB, JS, DB and JAM designed experiments; XZ, GG, PS, AM, HB, JMH, JH, EB, AK, SN, DM and PB generated data; XZ, GG, PS, AM, HB, JMH, JH, PB, DM, AK, JS, DB and JAM analysed and interpreted data; ST, HY and PACW provided essential reagents and expertise; XZ and JAM prepared the manuscript. All authors provided critical review of the manuscript.

## DATA AND MATERIALS AVAILABILITY

All data are available in the main text or the supplementary materials and the source data is available at Mendeley Repository. RNA sequencing data has been deposited at Gene Expression Omnibus (GEO), with accession number PRJNA991760.

