## Supplementary figures for "Hypoxia inducible factors inhibit respiratory syncytial virus infection by modulation of nucleolin expression"

### Hypoxic and pharmacological activation of HIFs inhibits respiratory syncytial virus infection

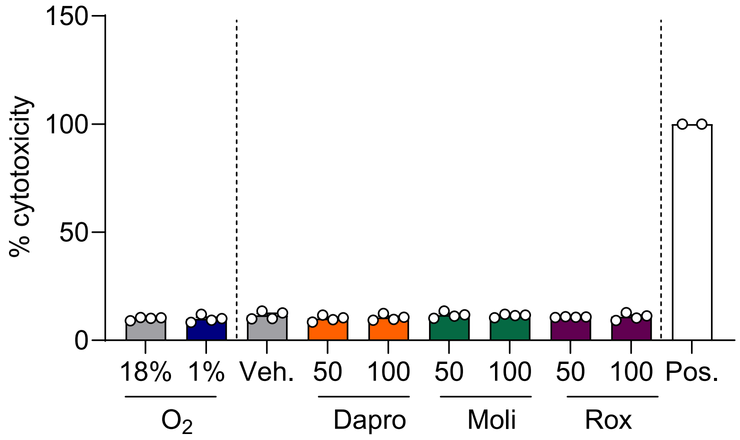


**Supplementary Figure 1. Cytotoxicity measurements of hypoxic or PHI treated Calu-3 cells.** Calu-3 cells were cultured at 18% or 1% % O2 or treated with PHIs (50 µM or 100 µM) for 72 h and cytotoxicity determined using an LDH assay (mean ± SEM, *n* = 4). Data are expressed relative to the positive control representing total cell lysate (100% cytotoxicity).

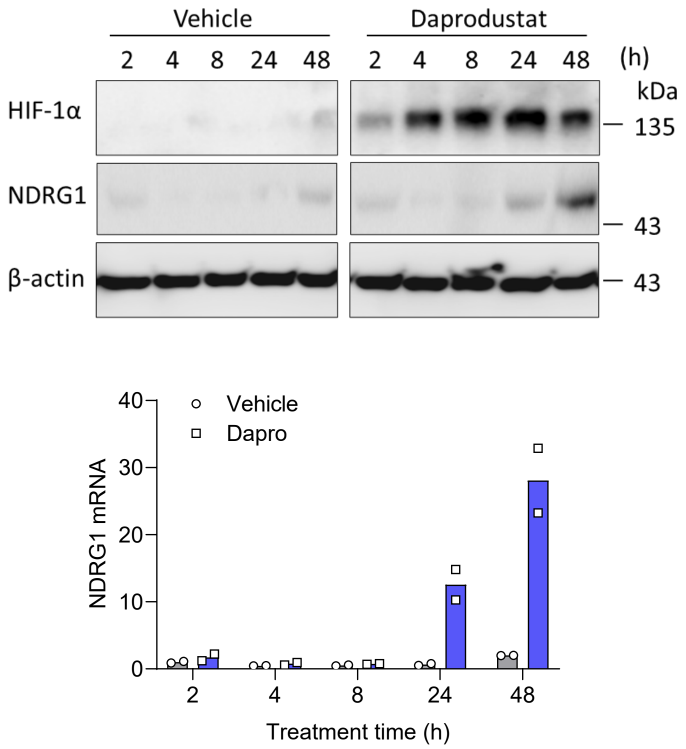


**Supplementary Figure 2.** Calu-3 cells were treated with Daprodustat at 50 µM and samples collected at 2, 4, 8, 24 or 48h. HIF-1α, NDRG-1 and β-actin protein expression were assessed by immunoblot. NDRG-1 mRNA was determined by qPCR and data expressed relative to the untreated cells at 2 h.

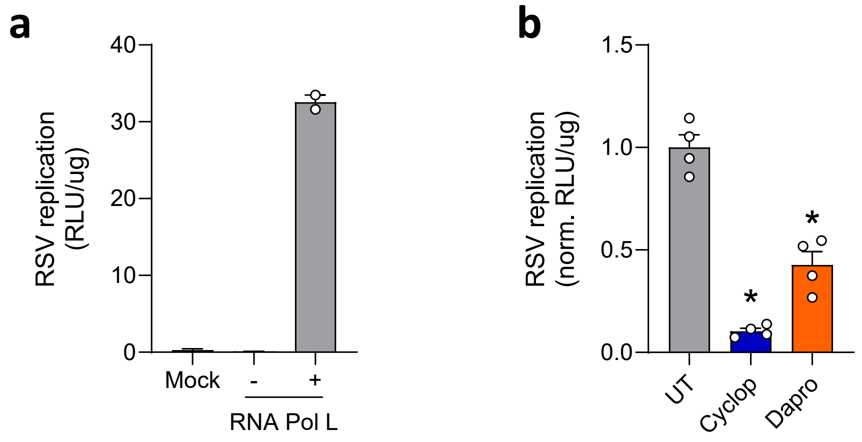


**Supplementary Figure 3. Inhibition of RSV replication by Daprodustat**. **a.** Calu-3 cells expressing T7 were transfected with plasmids encoding the RSV replicase or 'RSV replicon' with or without RNA polymerase L. Where the mock results derived from untransfected (mean ± SEM, n = 2). **b**. Calu-3 cells were transfected with RSV replicon for 24 h followed by treatment with Cyclopamine (10 µM) or Daprodustat (50 µM) and viral replication measured after 48 h (mean ± SEM, n = 4, Mann–Whitney test, Two-sided). Where UT denotes untreated RSV replicon bearing cells.

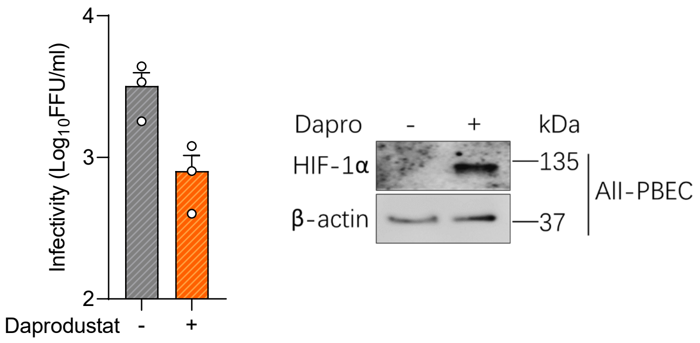


**Supplementary Figure 4. RSV infection is suppressed by treatment with Daprodustat.** Air Liquid Interface primary bronchial epithelial (ALI-PBEC) cells were pre-treated with Daprodustat (100 µM) for 24 h before infecting with RSV (MOI 0.2). Infectivity of RSV secreted from the apical surface of the culture was measured at 48 hpi (mean ± SEM, n = 3). HIF-1α and β-actin protein expression were assessed by immunoblot at 48 hpi.

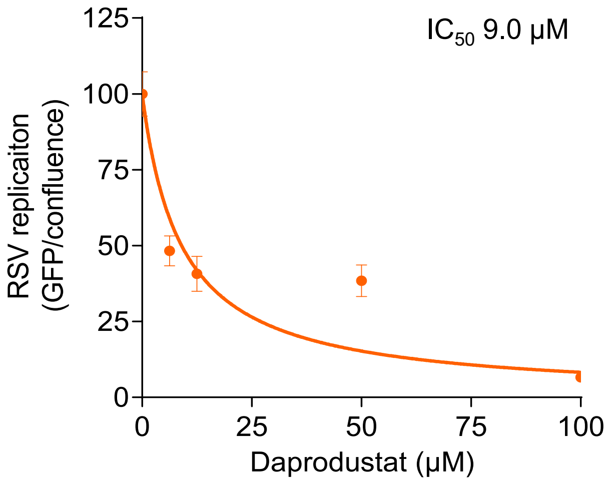


**Supplementary Figure 5. Dose-dependent inhibition of RSV infection by Daprodustat**. Calu-3 cells were pre-treated with a range of doses of Daprodustat for 24h prior to infection RSV-GFP (MOI 0.2) and viral replication determined over a 48 h period using an Incucyte® Live-Cell Imaging System. GFP signal was normalised per image and cell confluence assessed (mean ± SEM, n = 4, $r^2$ = 0.783).

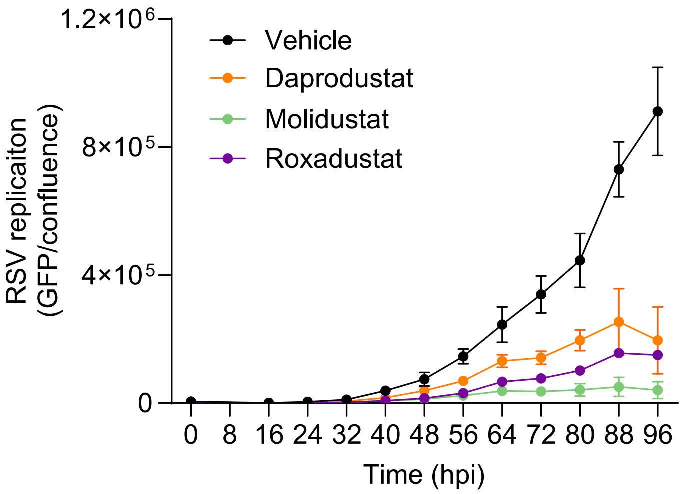


**Supplementary Figure 6. PHIs inhibit bovine RSV replication.** MDBK cells were pre-treated with Daprodustat, Molidustat, or Roxadustat for 24 h prior to infection with bovine RSV-GFP (MOI 0.2) and viral replication monitored at 8 h intervals using an Incucyte® Live-Cell Imaging System. GFP signal was normalized per image and cell confluence assessed (mean ± SEM, n = 4). Where Vehicle denotes untreated infected cells.

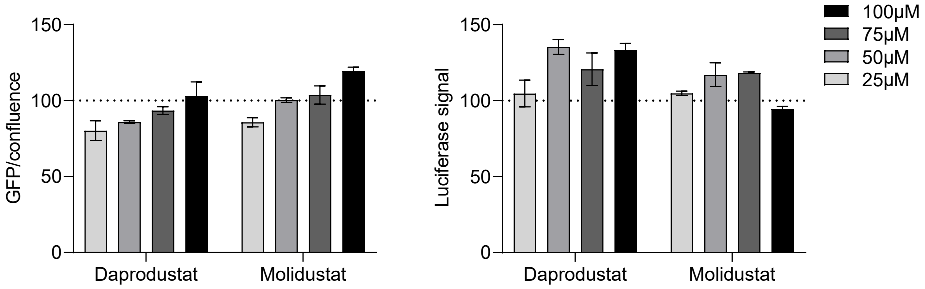


**Supplementary Figure 7. Null effect of Daprodustat or Molidustat on Fusion reporter activity.** HEK293T effector cells expressing beta strands 1-7 of GFP and RSV F protein were transfected with a plasmid encoding strands 8-11 of GFP and the complementary half of renilla luciferase. 24 h post transfection the cells were treated with different doses of PHIs in the presence of Doxycycline. After 48 h = the GFP signal was measured using the Incucyte system and luminescence measured by treating the cells with coelenterazine-H substrate. All values were normalised to the vehicle only samples (mean ± SEM, n = 2).

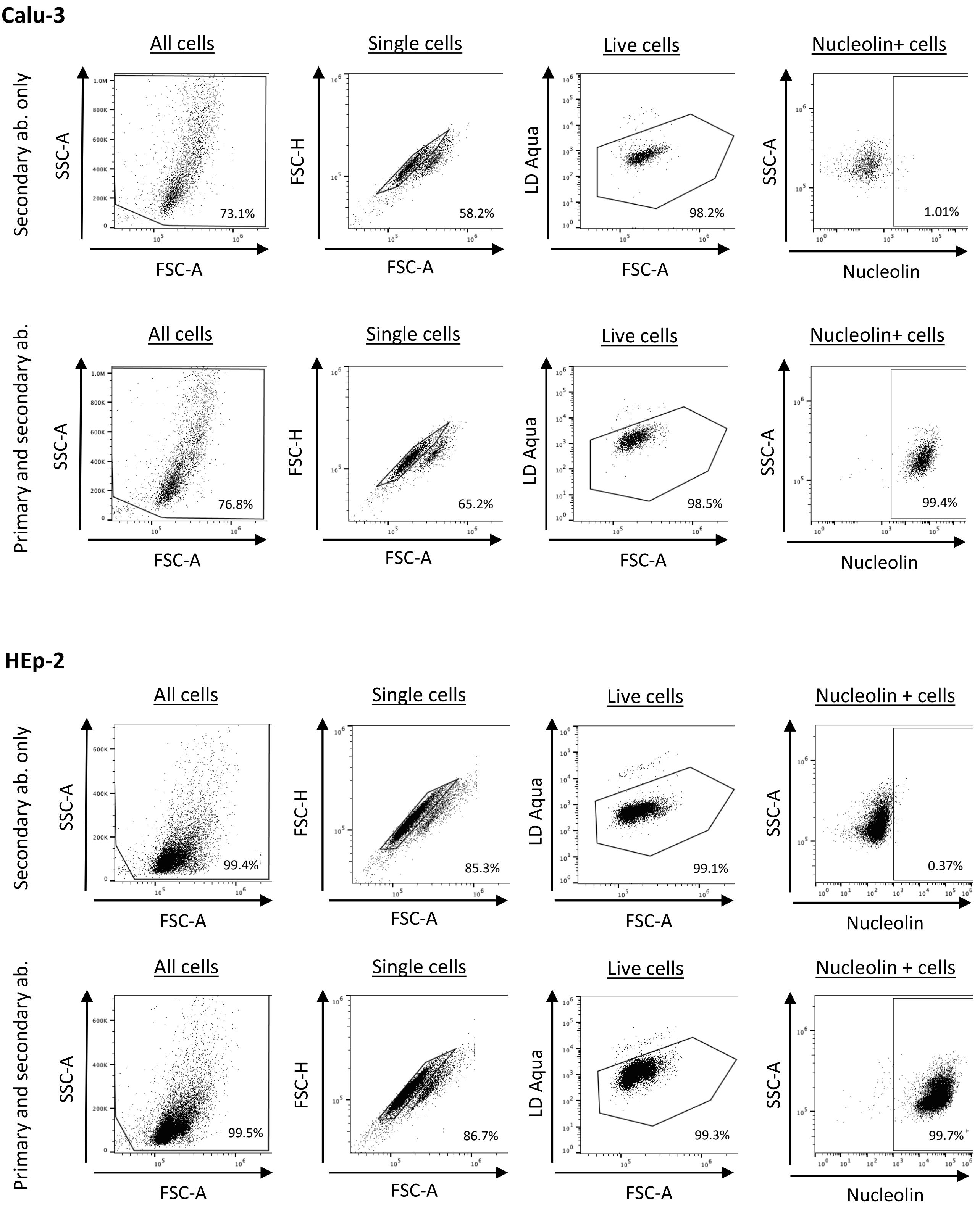


**Supplementary Figure 8. Flow cytometry gating strategy.** Representative dot plots illustrating gating strategy and analysis of untreated Calu-3 or HEp-2 cells which were stained with Live-Dead (LD) stain, and either the secondary antibody (goat anti-rabbit Alexa Fluor 488) only, or primary antibody (rabbit mAb to nucleolin) and secondary antibody. Gates were drawn based on the secondary only control. SSC = Side Scatter, FSC = Forward Scatter.

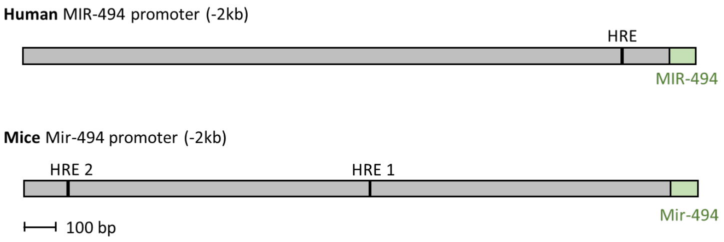


**Supplementary Figure 9. Location of the hypoxia response element (HRE) in the promoter region of human and mouse microRNA-494.** Base pairs (bp) are shown relative to the transcriptional start site.

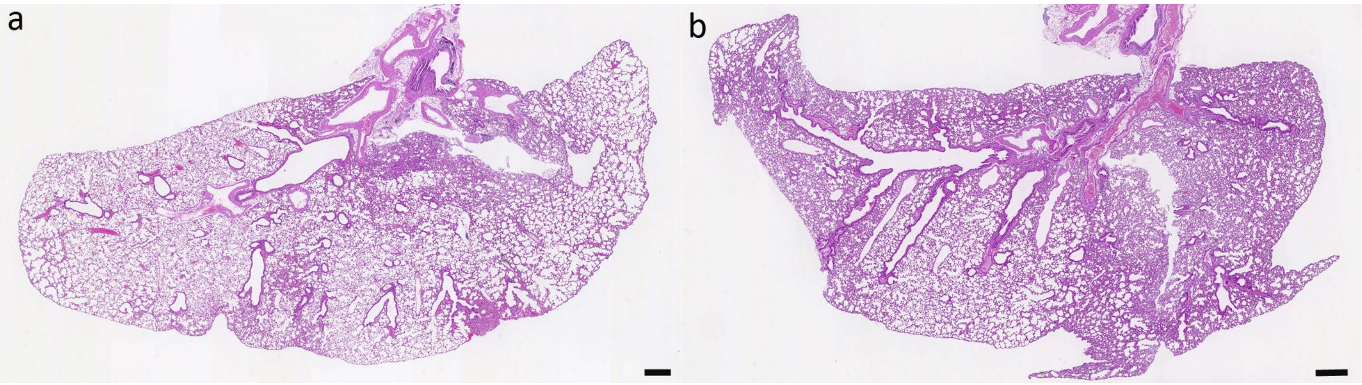


**Supplementary Figure 10. Daprodustat treatment does not have pathological effects in the lung.** Representative images of left lung lobes from vehicle-treated (**a**) or Daprodustat-treated (**b**) mice after 4 days. In both animals, the lung does not exhibit any histological changes. HE stain. Bars = 500 µm.

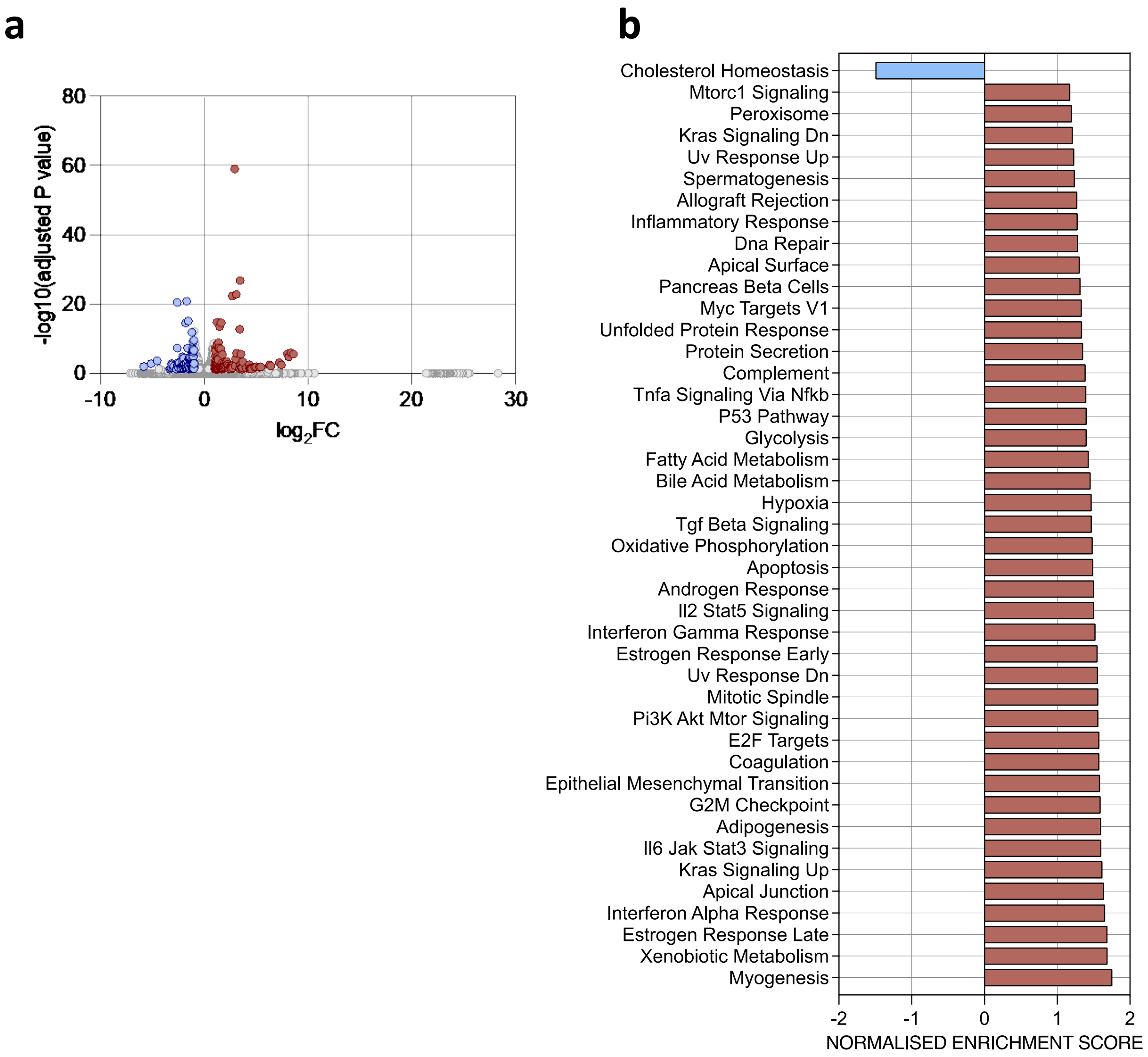


**Supplementary Figure 11. Transcriptomic analysis of the lung from vehicle or Daprodustat treated mice. a.** The impact of daprodustat treatment on the murine lung transcriptome. RNAs were extracted from the lung of Daprodustat (30mg/kg) treated and vehicle animals, and Illumina sequencing performed. DESEQ2 identified differential expressed genes, and a log2FC of +/- 1 and AdjP <0.05 was the threshold for statistical significance. **b**. Pathway analysis shows pertubation of host cell processes in response to Daprodustat treatment. Gene set enrichment analysis was performed using the Hallmarks gene sets from the moelcular signatures database. Gene sets with an FDR<0.25 are plotted, with red showing elevation in daprodustat treatment and blue representing pathways enriched in the vehicle.

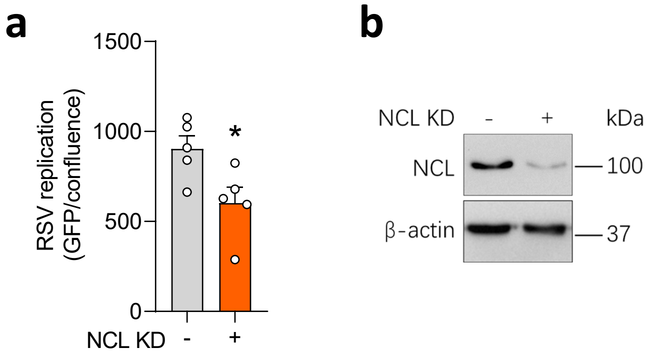


**Supplementary Figure 12. RSV infection is NCL dependent. a**. Calu-3 cells were transfected with siRNA targeting NCL 24h prior to infection with RSV-GPF (MOI 0.2) and viral replication measured using an Incucyte® Live-Cell Imaging System at 48hpi. GFP signal was normalized per image and cell confluence assessed (mean ± SEM, *n* = 5, Mann–Whitney test, Two-sided). **b**. Immunoblotting cell lysates for NCL expression following siRNA KD at 48hpi.
